# Dynamics of neuropeptide release and activity post-bloodmeal in the female mosquito, *Aedes aegypti*

**DOI:** 10.1101/2025.01.04.631332

**Authors:** Farwa Sajadi, Chiara Di Scipio, Lulia Snan, Jean-Paul Paluzzi

## Abstract

Female *Aedes aegypti* secrete urine rapidly post-bloodmeal ingestion, with diuresis beginning immediately for removal of excess salts and water. This post-prandial diuresis includes a peak, post-peak, and late phase, involving the combined actions of multiple hormones, including diuretic and anti-diuretic factors. Calcitonin-like diuretic hormone 31 (DH_31_) and kinin peptides stimulate diuresis through actions on their cognate receptors localized in the Malpighian ‘renal’ tubules (MTs). In contrast, the anti-diuretic neurohormone, CAPA, inhibits secretion by MTs stimulated by select diuretic hormones, including DH_31._ While DH_31_ and kinin are critical in achieving post-prandial diuresis, and CAPA functions as an important anti-diuretic hormone, the kinetics of their release and haemolymph levels remain unknown. Herein, using heterologously expressed *A. aegypti* DH_31_, CAPA, and kinin receptors, we investigated the titres of these hormones in the haemolymph of female mosquitoes at different time points after blood feeding. Haemolymph extracts from female mosquitoes contained levels of diuretic peptides, specifically kinin and DH_31_, that increased immediately post-bloodmeal, with levels peaking at 2 and 5 min, respectively, while DH_31_ levels remained elevated for 15 min. Comparatively, levels of CAPA peptides in the haemolymph steadily increase 15 min post-blood feeding, with levels peaking at 30 min. Synergistic actions were observed between DH_31_ and a kinin-like peptides on the MTs, providing a physiological context for the rapid release of these peptides into the female haemolymph. Altogether, these results demonstrate that DH_31_ and kinin are released immediately post-bloodmeal and, along with CAPA peptides, have a coordinative action on the MTs to maintain haemolymph homeostasis through regulation of primary urine secretion.

**Summary statement:** This study characterizes neuropeptidergic regulators of the Malpighian ‘renal’ tubules by quantifying their circulating titres and release post-blood feeding in the female mosquito, *A. aegypti*.

## 2. Introduction

Female *A. aegypti* mosquitoes ingest bloodmeals equivalent to more than twice their body mass resulting in an enormous payload to the flying animal, while also making them more susceptible to predation (Roitberg et al., 2003). The bloodmeal, imbibed with each reproductive cycle, serves as a source of protein, nutrients, and vitamins for egg development (Beyenbach and Petzel, 1987), but also contains considerable amounts of unwanted salts and water that threaten haemolymph homeostasis (Beyenbach, 2003). To counter this challenge, a rapid natriuresis and diuresis commences before the bloodmeal is completed, with almost half the imbibed plasma volume and salt excreted within 1-2 hours of feeding (Williams et al., 1983). In female *A. aegypti*, post-prandial diuresis is observed in three phases: peak phase (within 10 min of feeding), post-peak phase (10–50 min), and late phase (50 to beyond 120 min), with significant water eliminated during the peak phase (Williams et al., 1983). During these distinct post-prandial phases of diuresis, ion composition in the urine varies, with greatest natriuresis (Na^+^ secretion) occurring during the peak phase (Beyenbach, 2003), and kaliuresis (K^+^ secretion) delayed until post-peak and late phases. The varied pattern of water and salt loss indicates that multiple regulatory mechanisms act simultaneously during post-prandial diuresis (Williams et al., 1983).

Evidence from previous studies indicates that endocrine control of this post-prandial diuresis in mosquitoes is regulated by the actions of both diuretic and anti-diuretic hormones, including a calcitonin-like diuretic hormone 31 (DH_31_) (Coast et al., 2005; Kwon et al., 2012; Sajadi et al., 2018), a corticotropin-releasing factor (CRF)-like diuretic hormone 44 (DH_44_) (Clark and Bradley, 1996; Clark et al., 1998a; Clark et al., 1998b; Jagge and Pietrantonio, 2008; Sajadi et al., 2018), insect kinins (Lu et al., 2011; Sajadi et al., 2018; Schepel et al., 2010; Veenstra, 1994), biogenic amines such as 5-hydroxytryptamine (serotonin, 5HT) (Ionescu and Donini, 2012; Veenstra, 1988; Sajadi et al., 2018), and CAPA neuropeptides (Ionescu and Donini, 2012; Pollock et al., 2004; Sajadi et al., 2018; Sajadi et al., 2020). These endocrine messengers act on cognate receptors in the renal organs, the Malpighian tubules (MTs), to regulate ion and water transport for primary urine production (Beyenbach, 2003). In the *Aedes* mosquito, MTs are comprised of two primary cell types forming a simple epithelium; mitochondria-rich principal cells, which facilitate active cation (Na^+^ and K^+^) transport from the haemolymph into the tubule lumen, and thin less-abundant stellate cells that aid in the transepithelial secretion of Cl^−^ (Beyenbach, 2003; Patrick et al., 2006). Post-bloodmeal, excess fluid and ions (Na^+^, K^+^, and Cl^−^) are transported from the haemolymph into the tubule lumen, secreting high rates of urine (Beyenbach, 2003; Lu et al., 2011). Transport is achieved by channels and cotransporters in the principal and stellate cells, driven by the V-type H^+^-ATPase (VA) (Beyenbach, 2003; Patrick et al., 2006; Wieczorek et al., 1991) expressed in the apical brush border membrane of principal cells (Weng et al., 2003). The initial stimulus for this diuresis and natriuresis post-blood feeding is mosquito natriuretic peptide (MNP), which is released into the haemolymph when the female imbibes a bloodmeal (Beyenbach and Petzel, 1987). The calcitonin-like hormone, DH_31_, was identified as the MNP in the mosquitoes *A. aegypti* and *Anopheles gambiae* (Coast et al., 2005), activating transepithelial secretion of Na^+^ in the MTs via the second messenger, cyclic AMP (cAMP) (Beyenbach, 2003). Upon release from the CNS, DH_31_ binds to a G protein-coupled receptor (GPCR) localized in select principal cells with a distal-proximal gradient of expression (Kwon et al., 2012), increasing cAMP production (Beyenbach and Petzel, 1987), upregulating VA function and assembly that ultimately increases fluid secretion (Beyenbach, 2003; Karas et al., 2005; Sajadi et al., 2023). Another group of diuretic hormones are the insect kinin family, a group of multifunctional neuropeptides with both diuretic and myotropic actions (Nachman et al., 2009). There are three endogenous *A. aegypti* kinin peptides, encoded by a single cDNA, that bind to the kinin receptor localized within stellate cells, increasing levels of intracellular Ca^2+^ via 1,4,5-triphosphate (IP_3_) signaling to stimulate fluid secretion (Cady and Hagedorn, 1999a; Cady and Hagedorn, 1999b; Lu et al., 2011; Veenstra, 1994; Veenstra et al., 1997).

In adult female *A. aegypti* MTs, CAPA neuropeptides elicit an anti-diuretic role, with CAPA-1 shown to inhibit secretion by MTs stimulated by select diuretic factors, namely DH_31_ and 5HT, whilst having no effect on other diuretic factors, including kinin-related and DH_44_-related peptides (Sajadi et al., 2018). CAPA peptides bind to a GPCR localized exclusively to the principal cells (Sajadi et al., 2020), eliciting their actions through the nitric oxide synthase (NOS)/ cyclic GMP (cGMP)/ protein kinase G (PKG) pathway (Ionescu and Donini, 2012; Pollock et al., 2004; Terhzaz et al., 2012; Sajadi et al., 2020) leading to reduced VA activity and assembly, which halts fluid secretion by the MTs (Sajadi et al., 2023).

While haemolymph levels of DH_31_, kinin, and CAPA have yet to be determined, earlier studies suggested that DH_31_ is immediately released into circulation post-bloodmeal, allowing for the rapid secretion of Na^+^ and excess water (Coast et al., 2005). Additionally, kinin peptides have been proposed to be active during the peak phase of diuresis since knockdown of the kinin receptor reduces the rate and total volume of urine excreted (Kersch and Pietrantonio, 2011). In the present study, the *A. aegypti* DH_31_, kinin, and CAPA receptors identified previously (Kwon et al., 2012; Lu et al., 2011; Sajadi et al., 2020) were heterologously expressed to determine the circulating titres of DH_31_, kinin, and CAPA peptides in the haemolymph of blood fed females to determine the kinetics of release of both diuretic and anti-diuretic hormones in the adult female mosquito. Using this approach, we found that kinin and DH_31_ peptides are released into the female haemolymph almost instantaneously, peaking at about 2 and 5 minutes post-bloodmeal, respectively. In contrast, CAPA neuropeptides were found to rise more slowly in the haemolymph after blood-feeding, peaking at 30 minutes post-bloodmeal. In light of these new findings confirming the presence of multiple hormones regulating the MTs, we wondered whether they would affect fluid secretion rates in an additive or synergistic manner. Indeed, *in vitro* application of both DH_31_ and a kinin-like peptide revealed that these diuretics act synergistically in stimulating fluid secretion, providing novel evidence of coordinated action between DH_31_ and kinin released post-blood feeding. This rise in both diuretic and anti-diuretic levels correspond with the immediate and short-lived peak phase of diuresis after a female mosquito ingests a bloodmeal, further uncovering the precise regulatory mechanisms acting to clear excess ions and water to reduce the potential of predation, while simultaneously preventing disturbances to the hydromineral balance of their haemolymph, which ultimately contributes towards the success of this notorious human-disease vector.

## 3. Materials and methods

### 3.1 Animals

*Aedes aegypti* (Liverpool) eggs were collected from an established laboratory colony, and hatched in double-distilled water in an incubator at 26°C on a 12:12 hour light:dark cycle. Larvae were fed a solution of 2% (w/v) brewer’s yeast, 2% (w/v) Argentine beef liver powder (NOW foods, Bloomingdale, IL, USA), and adults were provided with a 10% sucrose solution *ad libidum*. Colony upkeep involved feeding adult females with sheep’s blood in Alsever’s solution weekly (Cedarlane Laboratories Ltd., Burlington, ON, Canada) using an artificial feeding system described previously (Rocco et al., 2017). All experiments were performed on four to seven-day old females that were either sucrose-fed or blood-fed as outlined in each specific experiment.

### 3.2 Haemolymph collection

To collect haemolymph samples, adult *A. aegypti* female mosquitoes were divided in two treatment conditions; 1) non-blood-fed (control) females provided 10% sucrose solution *ad libidum* and 2) blood-fed females also provided 10% sucrose solution *ad libidum*. Females in the blood-fed condition were fed to repletion and subsequently isolated after 0, 2, 5, 10, 15, 30, 60, 90 and 120 min post-bloodmeal (time = 0 min corresponds to the time immediately post-bloodmeal). Each female mosquito (either blood fed or control) was carefully opened at the segmental line between the last two abdominal segments within a 70 µL droplet of nuclease- and protease-free Dulbecco’s phosphate-buffered saline (DPBS; Wisent Corp., St. Bruno, QC, Canada) and haemolymph was allowed to diffuse into the DPBS droplet, all while ensuring no organs or blood-filled midguts ruptured. The surgically opened females were incubated in the DPBS droplet for 10 min, and the haemolymph DPBS mixture was subsequently collected and stored at −20°C.

### 3.3 Preparation of mammalian expression constructs

Gene-specific primers were designed to amplify the open reading frame of the *Aedae*DH_31_ receptor and *Aedae*Kinin receptor based on previously published sequences (Genbank Accession Numbers: DH_31_-R, AAGE02017873 (Kwon et al., 2012); Kinin-R, AY596453.1 (Lu et al., 2011); see Table 1). The amplified PCR products were purified using a Monarch PCR purification kit (New England Biolabs, Whitby, ON, Canada) and reamplified using modified primers containing restriction sites for subsequent directional cloning in mammalian expression plasmids. Products were A-tailed and cloned into a pGEM-T Easy vector (Promega, Madison, WI, USA) or pCE2 TA/Blunt-Zero vector (GeneBio Systems, Burlington, ON, Canada) and then transformed into 5-alpha high efficiency competent *E. coli* (New England Biolabs, Whitby, ON, Canada). Base accuracy of purified plasmid DNA miniprep was confirmed by Sanger sequencing (The Centre for Applied Genomics, Sick Kids, Toronto, ON, Canada). The receptor constructs DH_31_-R and Kinin-R were excised using standard restriction enzyme digestion and subcloned into the mammalian expression vectors, pBudCE4.1 Gα15 or pcDNA3.1^+^ (Life Technologies, Burlington, ON, Canada), with the former containing a mammalian promiscuous G protein (Gα15), which facilitates coupling of a wide variety of GPCRs to calcium signaling (Wahedi et al., 2019). The previously characterized *A. aegypti* CAPA receptor inserted into pcDNA3.1^+^ (Sajadi et al., 2020) was also used for transfection of mammalian cells. Plasmid DNA was purified from overnight bacterial cultures using the Monarch Plasmid Miniprep Kit (New England Biolabs, Whitby, ON, Canada) following manufacturer guidelines. These three heterologously expressed receptors were used to determine the timing of release and quantify the levels in the haemolymph of CAPA (both CAPA-1 and CAPA-2), DH_31_ along with kinin neuropeptides in association with blood feeding by adult female mosquitoes.

**Table 1.**
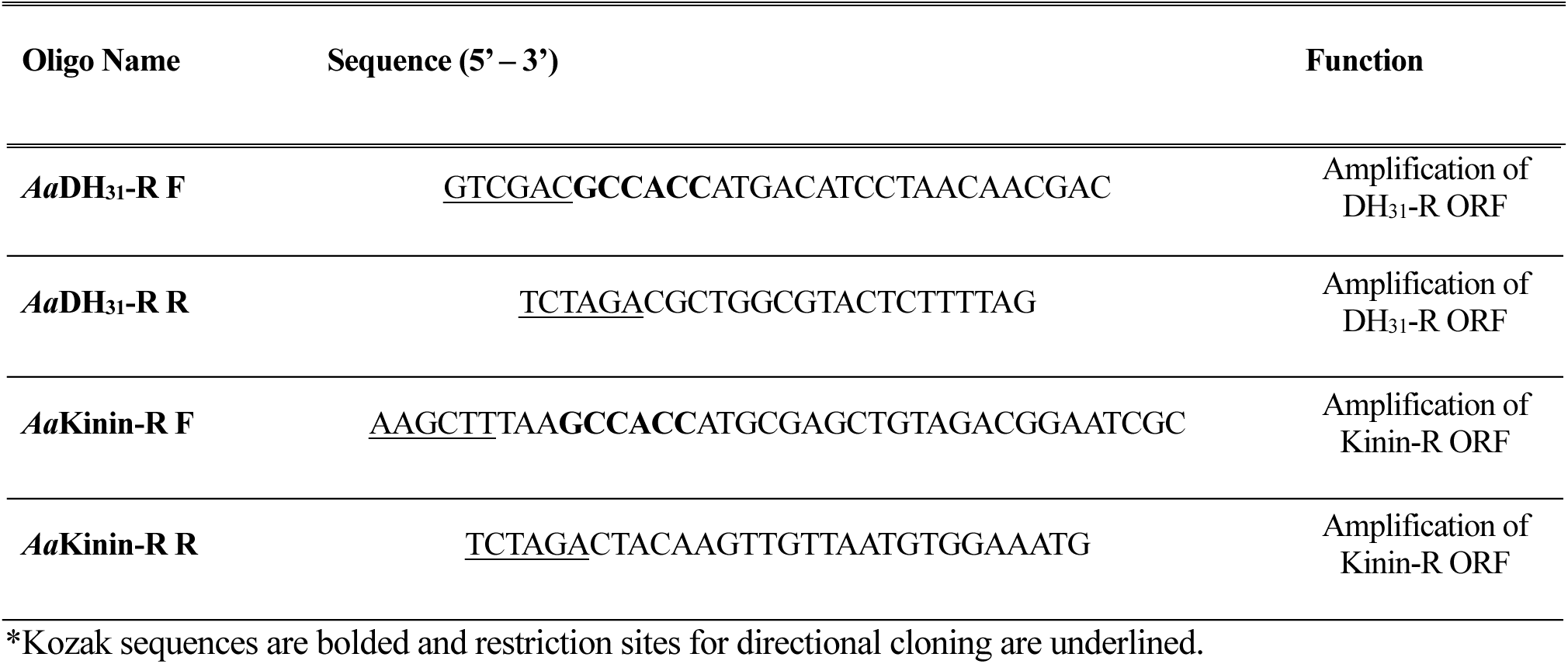
Gene-specific primer information used for amplification of the complete cDNA of *A. aegypti*.

### 3.4 Heterologous receptor functional activation bioluminescence assay

Functional activation of the *Aedae*DH_31_-R, *Aedae*CAPA-R, and *Aedae*Kinin-R was assayed using a previously established recombinant Chinese hamster ovary (CHO)-K1 cell line stably expressing aequorin (Paluzzi et al., 2012a; Wahedi and Paluzzi, 2018). Cells were grown and maintained in Dulbecco’s modified eagles medium:nutrient F12 (DMEM:F12; 1:1) media containing 10% heat-inactivated fetal bovine serum (FBS; Wisent, St. Bruno, QC, Canada), 1x antimycotic-antibiotic, and 200 µg/mL geneticin (G418 Sulfate). Mammalian cell lines were transiently transfected with either pBudCE4.1 Gα15 expression vector possessing *Aedae*DH_31_-R, pcDNA3.1^+^ expression vector possessing *Aedae*CAPA-R, or with pcDNA3.1^+^ expression vector possessing *Aedae*Kinin-R using Lipofectamine LTX and Plus Reagent transfection system (Invitrogen, Burlington, ON, Canada) following a 3:1:1 transfection reagent (µL): Plus reagent (µL): plasmid DNA (µg) as described previously (Wahedi and Paluzzi, 2018). Approximately 48h post-transfection, cells were dislodged from the culture flasks using 5 mmol l^−1^ ethylenediaminetetraacetic acid (EDTA; Life Technologies, Burlington, ON, Canada) in DPBS and cells were resuspended in BSA medium (DMEM-F12 media containing 0.1% bovine serum albumin, 1X antimycotic-antibiotic) at a concentration of 10^6^-10^7^ cells/mL quantified using a Countess II FL cell counter (ThermoFisher Scientific, Burlington, ON, Canada). Coelenterazine *h* (Promega, Madison, WI, USA) was added to the cells to a final concentration of 5 µM and incubated for 3 hours as described previously (Oryan et al., 2018), and subsequently, diluted 10-fold in BSA medium and incubated for an additional hour at room temperature. To measure *in vivo* circulating levels of DH_31_, kinin, and CAPA in female mosquitoes, collected haemolymph samples along with standards (synthetic peptides, see below) were loaded into 96-well white luminescence plates (Greiner Bio-One, Germany), and cells expressing either *Aedae*DH_31_-R, *Aedae*Kinin-R, or *Aedae*CAPA-R were loaded into each well with an automated injector unit and luminescence was immediately measured over 20 seconds using a Synergy 2 Multi-Mode Microplate Reader (BioTek, Winooski, VT, USA). To confirm receptor specificity, serial dilutions of various synthetic peptide ligands (purity >90%, Genscript, Piscataway, NJ, USA) were prepared in DPBS and loaded into quadruplicates in white 96-well luminescence plates. The various peptides used in this study are listed in Table 2. To test receptor specificity for the Kinin-R, two kinin-like peptides were used; *Culex* depolarizing peptide (CDP) and *Drosophila* leucokinin (LK) *in lieu* of *Aedes* kinins which we did not have available. Negative controls were carried out using DPBS alone while 5 µM ATP was used as a positive control, which activates endogenously expressed purinoceptors (Iredale and Hill, 1993; Michel et al., 1998). Experimental sample luminescent responses were normalized to ATP-induced maximal luminescent responses, multiplied by the dilution factor, and neuropeptide concentration levels in the haemolymph were determined through non-linear regression analysis in GraphPad Prism 9.1 (GraphPad Software, San Diego, CA, USA).

**Table 2.**
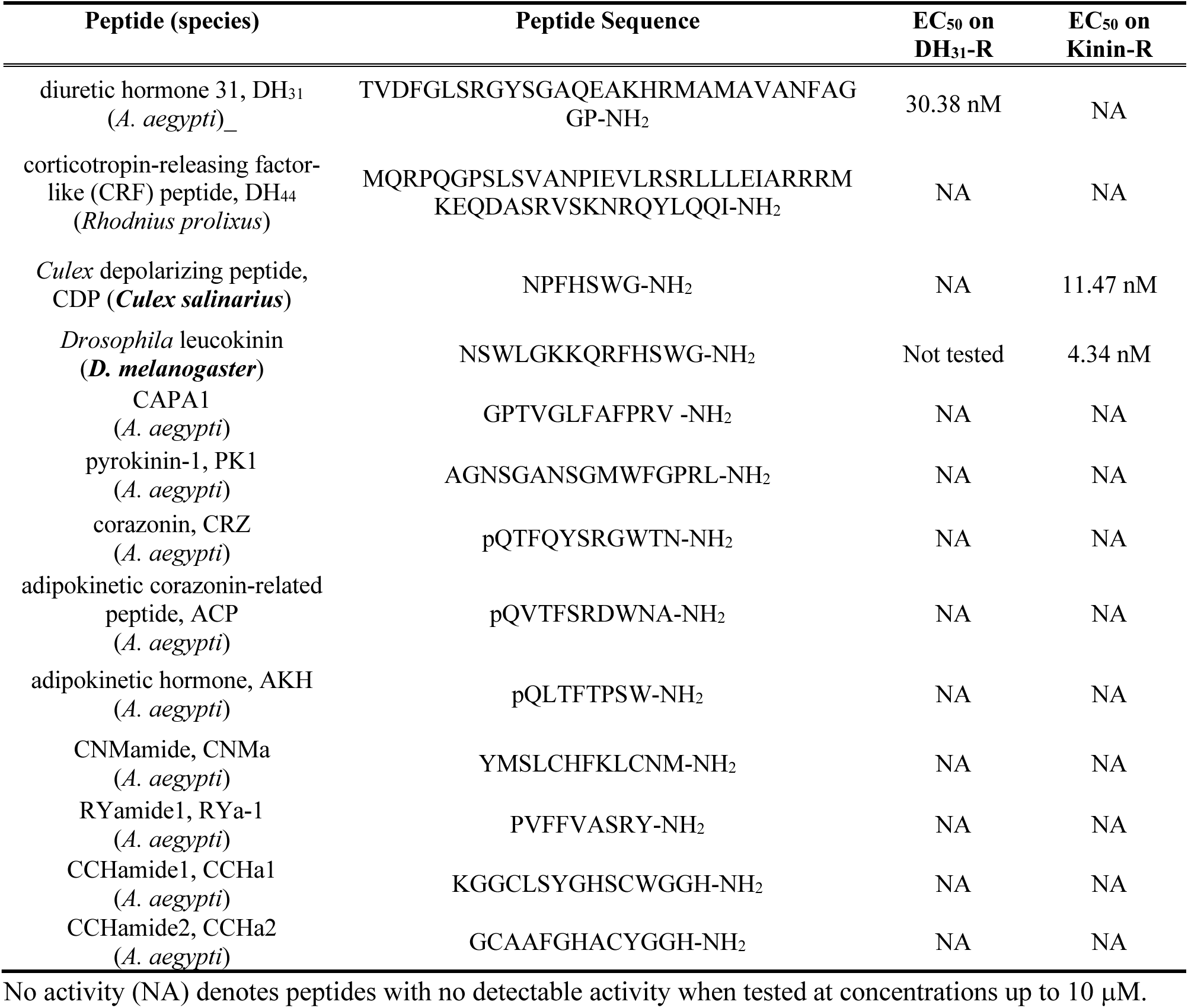
Name and sequence of *A. aegypti* (or other insect, as specified) neuropeptides used in the heterologous functional assays validating specificity of *Aedae*DH_31_-R and *Aedae*Kinin-R along with EC_50_ values.

### 3.5 Fluid secretion assay on individually isolated MTs

In order to determine fluid secretion rates, modified Ramsay assays were performed as described previously (Sajadi et al., 2021; Sajadi et al., 2018). Female adults (between 4-7 day old) were dissected under physiological *Aedes* saline adapted from (Petzel et al., 1987) that contained (in mmol^−1^): 150 NaCl, 25 HEPES, 3.4 KCl, 7.5 NaOH, 1.8 NaHCO_3_, 1 MgSO_4_, 1.7 CaCl_2_, and 5 glucose, and titrated to pH 7.1. Individual MTs were removed and placed in a Sylgard-lined Petri dish containing 20 µL bathing droplets (1:1 mixture of Schneider’s Insect Medium (Sigma-Aldrich, Oakville, ON, Canada): *Aedes* saline), immersed in hydrated mineral oil to prevent evaporation. The proximal end of each tubule was wrapped around a minutien pin to allow for fluid secretion measurements. Dose response analysis for *Aedae*DH_31_ was determined by using concentrations ranging from 10^−5^ to 10^−12^ mol l^−1^ on unstimulated MTs. To investigate the effects of collected haemolymph samples on fluid secretion rate, haemolymph samples from both non-blood fed and blood fed females were used against unstimulated MTs and allowed to incubate for 60 min before measurement of secreted fluid droplets. To investigate potential additive or synergistic activity, dosages of 25 nmol l^−1^ *Aedae*DH_31_, 10 nmol l^−1^ *Rhodnius prolixus* DH (CRF-related peptide, DH_44_), 50 nmol l^−1^ CDP, 100 nmol l^−1^ 5HT, and 1 fmol l^−1^ *Aedae*CAPA-1 were used (Sajadi et al., 2018; Sajadi et al., 2020).

### 3.6 Statistical analyses

Data was compiled using Microsoft Excel and transferred to Graphpad Prism software v.9 to create figures and conduct all statistical analyses. Data was analyzed accordingly using a one-way ANOVA and the appropriate post-hoc test for multiple comparisons or an unpaired t-test to determine whether there is a difference between two samples, as indicated in the figure captions, with differences between treatments considered significant if p<0.05.

## 4 Results

### 4.1 Functional characterization of AedaeDH_31_-R and DH_31_ hormone detection in haemolymph post-bloodmeal

A heterologous receptor functional assay involving CHO-K1 cells stably expressing a bioluminescent calcium sensor, aequorin (Paluzzi et al., 2012a; Wahedi and Paluzzi, 2018), was used to functionally characterize the *Aedes* DH_31_ receptor (*Aedae*DH_31_-R) identified earlier (Kwon et al., 2012). Our results confirm the specificity of the *Aedae*DH_31_-R for the *Aedae*DH_31_ peptide as demonstrated by the dose-dependent luminescent response (Fig. 1A) with a half maximal effective concentration (EC_50_) of 30.38nM. The activity of other *A. aegypti* peptides tested along with peptides having structural similarity to endogenous mosquito neuropeptides, including *Rhodnius prolixus* corticotropin-releasing factor (CRF)-like peptide (DH_44_), yielded no detectable activity over background luminescence levels (Fig. 1B, Table 2). Moreover, no luminescence signals above background were obtained in control transfections of CHO-K1-aeq cells expressing enhanced green fluorescent protein (EGFP) (data not shown) to any peptide used in this study, confirming the calcium-based luminescence response observed was a result of the DH_31_ neuropeptide activating the heterologously expressed *A. aegypti* DH_31_ receptor.

**Figure 1.**
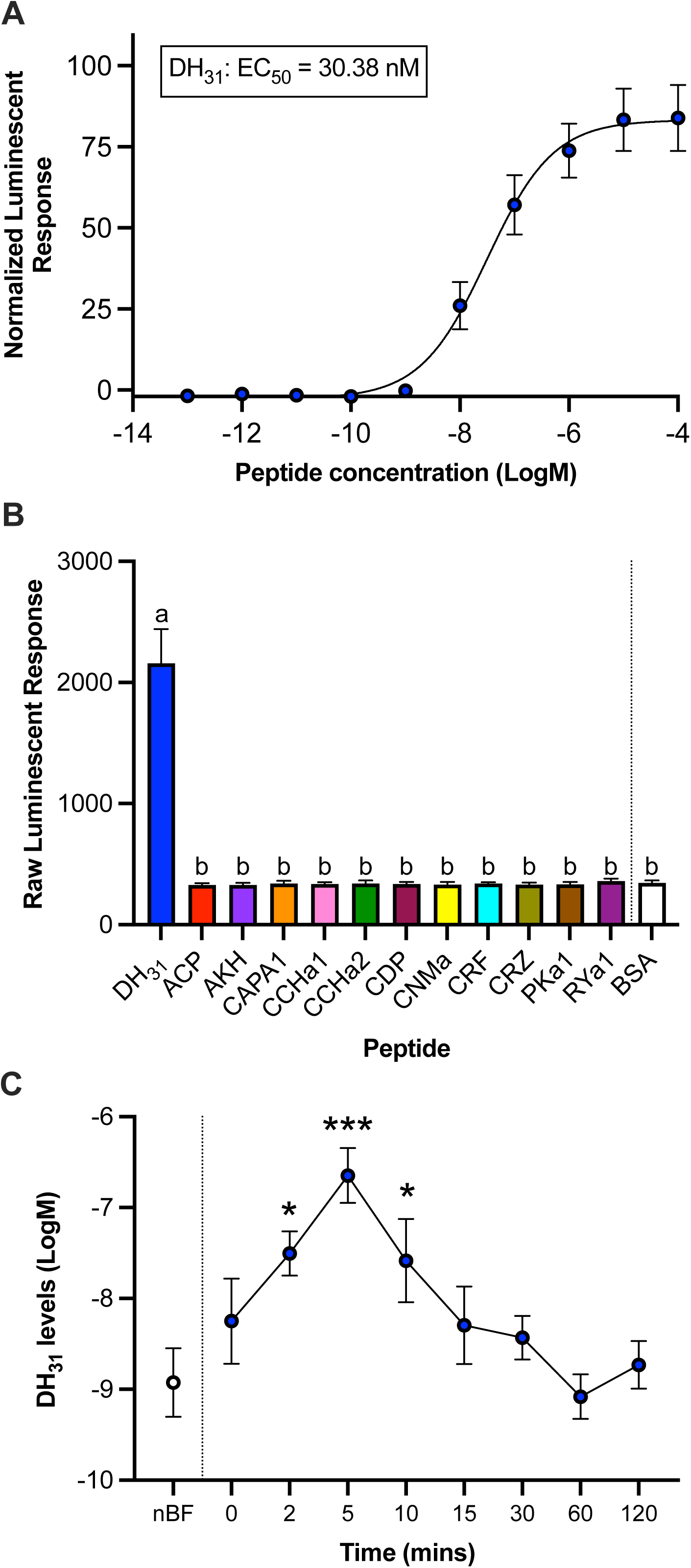
Functional heterologous receptor assay involving the *A. aegypti* DH_31_ receptor to determine *Aedae*DH_31_ titres in the haemolymph of blood fed female *A. aegypti*. (A) Normalized dose-response curve after the addition of 10^−13^ – 10^−4^ M doses of *Aedes*DH_31_ peptide. Luminescence was normalized to the BSA control and plotted relative to the maximal response (10^−4^ M). The EC_50_ for *Aedae*DH_31_ is 30.38 nM. (B) Raw luminescent response (average over 20 sec) following addition of 10^−6^ M dose of representative neuropeptides belonging to several insect peptide families (n=3); different letters denote bars that are significantly different from control (BSA assay media) as determined by a one-way ANOVA and Dunnett’s multiple comparison post-hoc test (p<0.0001). For peptide sequence information, see Table 2. (C) Females were blood fed and their haemolymph was collected post-feeding (see methods for details) and compared to non-blood fed (sugar-fed only) females. Endogenous concentrations of DH_31_ in female *A. aegypti* haemolymph following a bloodmeal and in non-blood fed (nBF) mosquitoes were determined by comparing to the dose-response curve using synthetic *Aedes*DH_31_ peptide. Significantly different DH_31_ levels in the haemolymph compared to nBF are denoted by asterisks as determined by a one-way ANOVA and Dunnett’s multiple comparison post-hoc test (* denotes p < 0.05 and *** denotes p < 0.0001); data represent the mean±SEM (n=7–12).

To examine the timing of release of DH_31_ in the *Aedes* mosquito, females were allowed to blood feed, and their haemolymph was subsequently collected after 0, 2, 5, 10, 15, 30, 60, and 120 min post-blood feeding. Using the heterologous receptor assay, we expressed the *Aedae*DH_31_-R and its activation was monitored following treatment with collected haemolymph samples, which allowed us to determine the circulating DH_31_ titres in the haemolymph by interpolation from the dose-response curve established using synthetic DH_31_ (Fig. 1C). Immediately post-blood feeding, DH_31_ levels in the female mosquito were determined to be 5.6±16.6 nM, which is comparable to non-blood-fed females (1.2±2.8 nM). Levels of DH_31_ increase significantly at 2 min after blood-feeding reaching 31.3±54.8 nM, peaking at 5 min to 224.9±450.6 nM and remain significantly elevated 10 min after blood-feeding at 26.1±74.8 nM (Fig. 1C). Thereafter, DH_31_ levels are reduced by 15 min and remain low up to 120 min after blood feeding, with haemolymph titres similar to those in non-blood fed females ranging between 0.8±1.5 nM to 5.1±13.6 nM. Considering the selectivity and specificity of the *Aedae*DH_31_-R for DH_31_ (Fig. 1A-B), this indicates that the luminescent response observed when heterologously expressed *Aedae*DH_31_-R was exposed to the haemolymph samples was a result of its activation by the endogenous DH_31_ peptidergic hormone.

### 4.2 CAPA peptides involved in inhibiting DH_31_-stimulated rapid diuresis post-bloodmeal

In order to characterize the timing and release of CAPA neuropeptides, CHO-K1 cells were transfected with the previously characterized anti-diuretic hormone receptor, *Aedae*CAPA-R (Sajadi et al., 2020). As a synthetic peptide standard used in the current study, the receptor was activated by the *Aedae*CAPA-1 peptide (EC_50_=1.83 nM), as demonstrated by the dose-dependent luminescent response (Fig. 2A) although the receptor is activated similarly by both endogenous CAPA neuropeptides, specifically *Aedae*CAPA-1 and *Aedae*CAPA-2 (Sajadi et al., 2020). Haemolymph samples collected from females at 0, 5, 10, 15, 30, 60, 90, and 120 min post-blood feeding were used for measuring levels of CAPA peptides using the heterologous assay. Normalized luminescent responses were interpolated from the dose-response curve generated with synthetic *Aedae*CAPA-1 peptide and circulating CAPA peptide hormone concentrations were determined (Fig. 2B). During non-blood fed conditions, haemolymph CAPA levels in females are at 72.4±113.7 pM. While CAPA concentrations had an increased trend over the first 15 minutes post-blood feeding, the levels did not reach statistical significance despite being over four-fold higher compared to non-blood fed mosquitoes. However, at 30 min after blood-feeding, the levels of CAPA within the haemolymph were measured at 931.1±1536.0 pM, which is significantly greater (by over twelve-fold) compared to CAPA peptide titres in the haemolymph of non-blood fed animals. The CAPA peptide levels in the haemolymph are reduced between 60 to 120 min, with levels indistinguishable from non-blood fed female mosquitoes. No luminescence signals greater than background were obtained when haemolymph samples were applied to cells transfected with empty pcDNA3.1^+^ vector (not shown), indicating the luminescent response observed when *Aedae*CAPA-R expressing cells were treated with the haemolymph samples was a result of the activation of this receptor by the endogenous CAPA peptide hormones.

**Figure 2.**
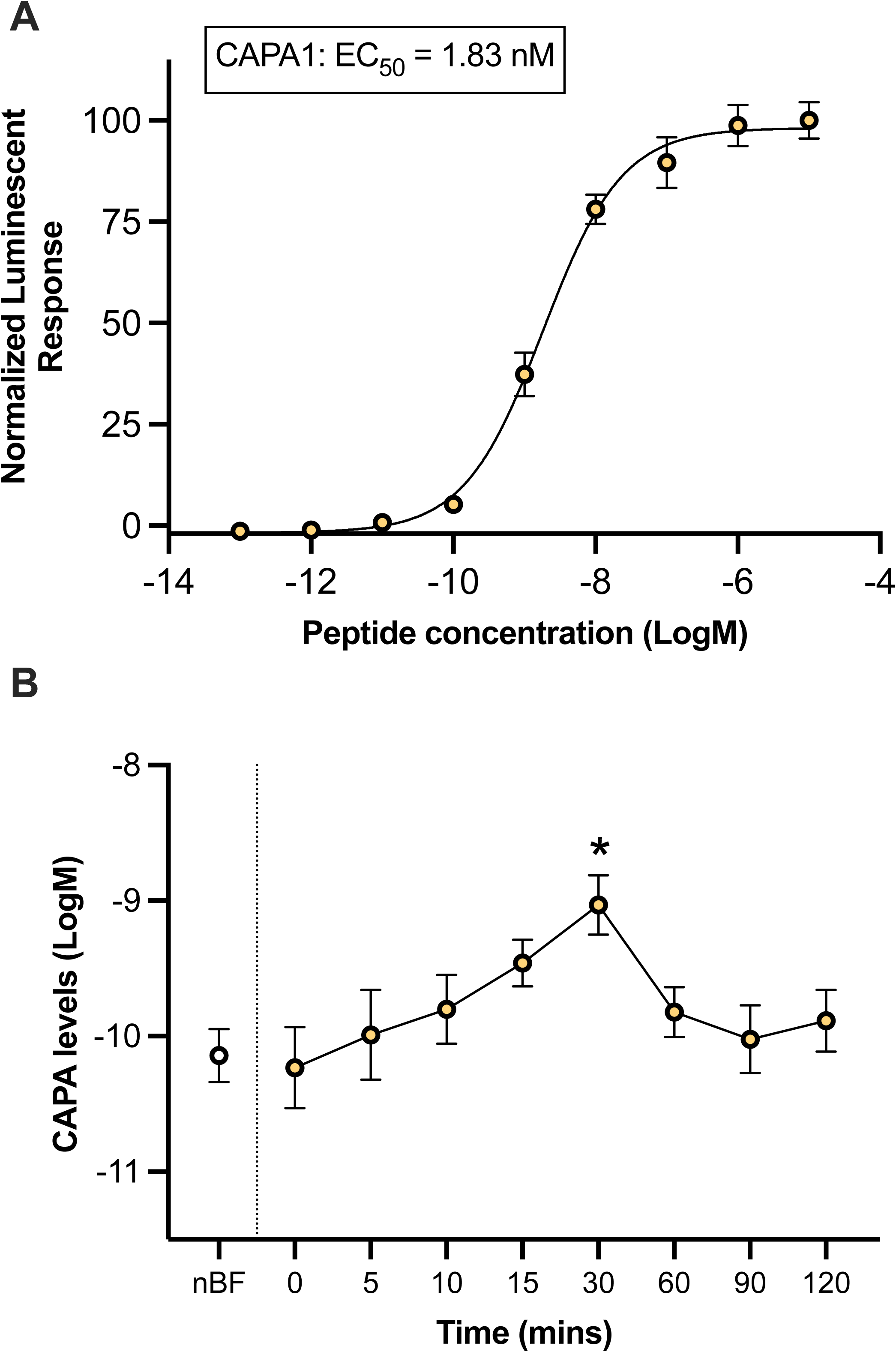
Functional heterologous receptor assay involving CHO-K1 cells expressing the *A. aegypti* CAPA receptor to determine CAPA titres in the haemolymph of blood fed female *A. aegypti*. (A) Normalized dose-response curve after the addition of 10^−13^ – 10^−5^ M doses of *Aedes*CAPA-1 peptide (EC_50_ = 1.83 nM). Luminescence was normalized to BSA media and plotted relative to the maximal response (10^−5^ M); data represent the mean±SEM (n=4). (B) Measured concentrations of CAPA peptide levels in female *A. aegypti* haemolymph following a bloodmeal (see methods for details on haemolymph collection, nBF = non-blood fed). In (B), significantly different CAPA peptide levels in the haemolymph compared to nBF are denoted by asterisks as determined by a one-way ANOVA and Dunnett’s multiple comparison post-hoc test (* denotes p < 0.05); data represent the mean±SEM (n=8–11).

### 4.3 Kinin peptides involved in immediate rapid diuresis post-bloodmeal

To examine the potential involvement of kinin in post-blood feeding diuresis, a similar heterologous receptor functional assay was used to quantify kinin peptide levels in the female *A. aegypti* haemolymph using the previously molecularly characterized *Aedae*Kinin-R (Lu et al., 2011). As a first step, the ligand specificity of *Aedae*Kinin-R had to be determined since no such data was available in the literature. Using CHO-K1 cells expressing aequorin, our results confirm that *Aedae*Kinin-R is activated specifically by kinin-like peptides including, CDP (EC_50_=11.47 nM) and LK (EC_50_=4.34 nM) (Fig. 3A). Specifically, no other tested endogenous mosquito neuropeptides or *Rhopr*DH_44_ (a corticotropin-releasing factor (CRF)-like diuretic peptide) displayed any detectable activity over background luminescence levels (Fig. 3B, Table 2). Haemolymph samples collected from females at different time points (similar to collections for quantifying DH_31_ levels) post-blood feeding were tested for presence of kinin peptides using the heterologous assay. Normalized luminescent responses were interpolated from the synthetic LK dose-response curve and kinin peptide concentrations in the haemolymph were determined. During non-blood fed conditions, haemolymph kinin levels in females are at 0.31±0.16 nM and remain at a similar level (0.11±0.06 nM) immediately post-blood feeding (0 min). However, by 2 min after blood feeding, kinin levels increase significantly (by over fifteen-fold), peaking at 4.78±4.19 nM (Fig. 3C). Thereafter, kinin levels decrease to levels similar to non-blood fed females at 5 min (0.18±0.09 nM) and remain in the sub-nanomolar range between 10 to 120 min post-blood feeding (range of 0.22±0.12 to 0.94±0.20 nM), which were not significantly different from levels detected in haemolymph of non-blood fed females (Fig. 3C). No luminescent response above background was observed by cells transfected with empty pcDNA3.1^+^ vector (not shown) against the synthetic kinins (or any other peptide) tested in this study, supporting *Aedae*Kinin-R specificity to the insect kinin-related peptides.

**Figure 3.**
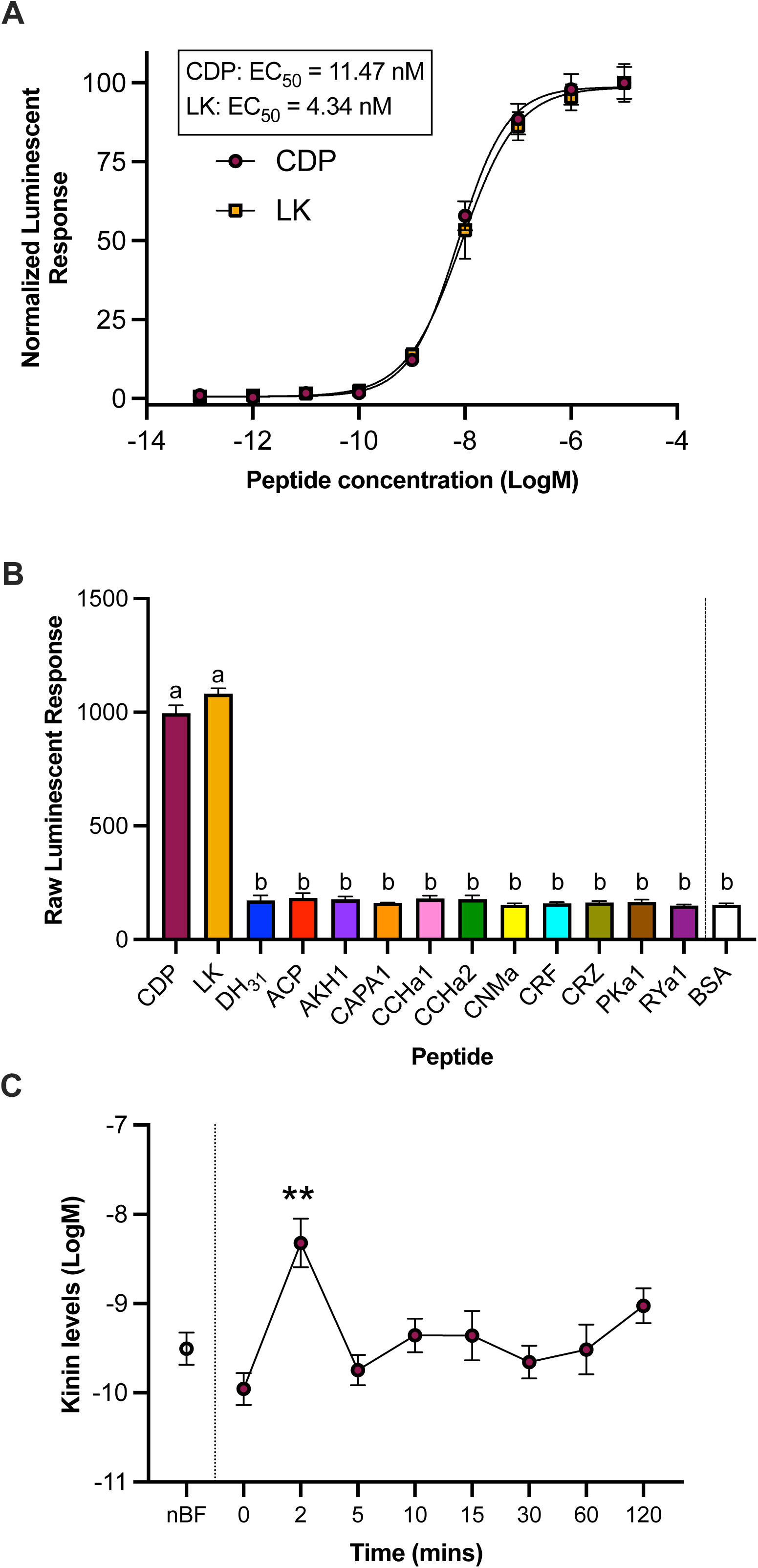
Functional heterologous receptor assay of CHO-K1 cells expressing the *A. aegypti* Kinin receptor to determine *Aedae*Kinin titres in the haemolymph of blood fed female *A. aegypti*. (A) Normalized dose-response curve after the addition of 10^−13^ – 10^−5^ M doses of CDP (EC_50_ = 11.47 nM) and Leucokinin (LK) (EC_50_ = 4.34 nM). (B) Raw luminescent response following addition of 10^−6^ M dose of representative neuropeptides belonging to several insect peptide families (n=3); different letters denote bars that are significantly different from control (BSA assay media) as determined by a one-way ANOVA and Dunnett’s multiple comparison post-hoc test (p<0.0001). For peptide sequence information, see Table 2. For peptide sequence information, see Table 2. (C) Concentrations of kinin peptides in female *A. aegypti* haemolymph following a bloodmeal (see methods for details on haemolymph collection, nBF = non-blood fed; relative to LK standard curve). Significantly different kinin peptide levels in the haemolymph compared to nBF are denoted by asterisks as determined by a one-way ANOVA and Dunnett’s multiple comparison post-hoc test (** denotes p < 0.001); data represent the mean±SEM (n=10–15).

### 4.4 Multiple hormones involved in regulating diuresis post-bloodmeal

To determine the *in vitro* potency of *Aedae*DH_31_ on isolated adult *A. aegypti* MTs, a wide range of dosages were applied against unstimulated tubules (Fig. 4A). Application of synthetic *Aedae*DH_31_ at doses ranging from 10^−5^ M to 10^−12^ M onto unstimulated adult tubules of *A. aegypti* resulted in a dose-dependent stimulation of fluid secretion with a half maximal response in the low nanomolar range (EC_50_=13.6 nM). Compared to basal secretion rates by unstimulated MTs, tested *Aedae*DH_31_ concentrations greater than 10nM (10^−8^ M) led to significantly higher secretion rates (Fig. 4A). To determine whether the collected haemolymph samples would also influence fluid secretion rates, and thereby reflect potential combined activity of peptides quantified using the heterologous assay, haemolymph samples were applied to MTs and incubated for 60 min (Fig. 4B). Haemolymph sample collected immediately post-blood feeding (0 min) caused a significant increase in fluid secretion (0.197±0.024 nL min^−1^) representing a four-fold increase in the secretion rate, in comparison to activity from non-blood fed haemolymph samples (0.048±0.016 nL min^−1^). Similarly high secretion rates were observed for MTs treated with haemolymph samples collected at 2 and 5 min post-feeding (0.203±0.024 nL min^−1^ and 0.174±0.027 nL min^−1^, respectively). Interestingly, haemolymph samples collected 10 – 30 min post-blood feeding demonstrated reduced stimulation of fluid secretion that was not significantly different from the activity of non-blood fed mosquito haemolymph (Fig. 4B). Following this trend, haemolymph samples collected from female mosquitoes between 60 to 120 min post-feeding resulted in secretion rates indistinguishable from non-blood fed females. Tubules treated with PBS alone, which was used as a diluent for the haemolymph sample collection, did not result in stimulation of fluid secretion rates. Rates of fluid secretion by MTs observed in response to haemolymph samples collected 0 to 5min post-blood-feeding were identical to those observed in response to synthetic *Aedae*DH_31_ (25 nmol l^−1^) whereas the activity of haemolymph samples collected 10 and 120 min post blood feeding was similar to MTs treated with *Aedae*DH_31_ (25 nmol l^−1^) combined with *Aedae*CAPA-1 (1 fmol l^−1^), which was indistinguishable to the activity of haemolymph from non-blood fed mosquitoes (Fig. 4B).

**Figure 4.**
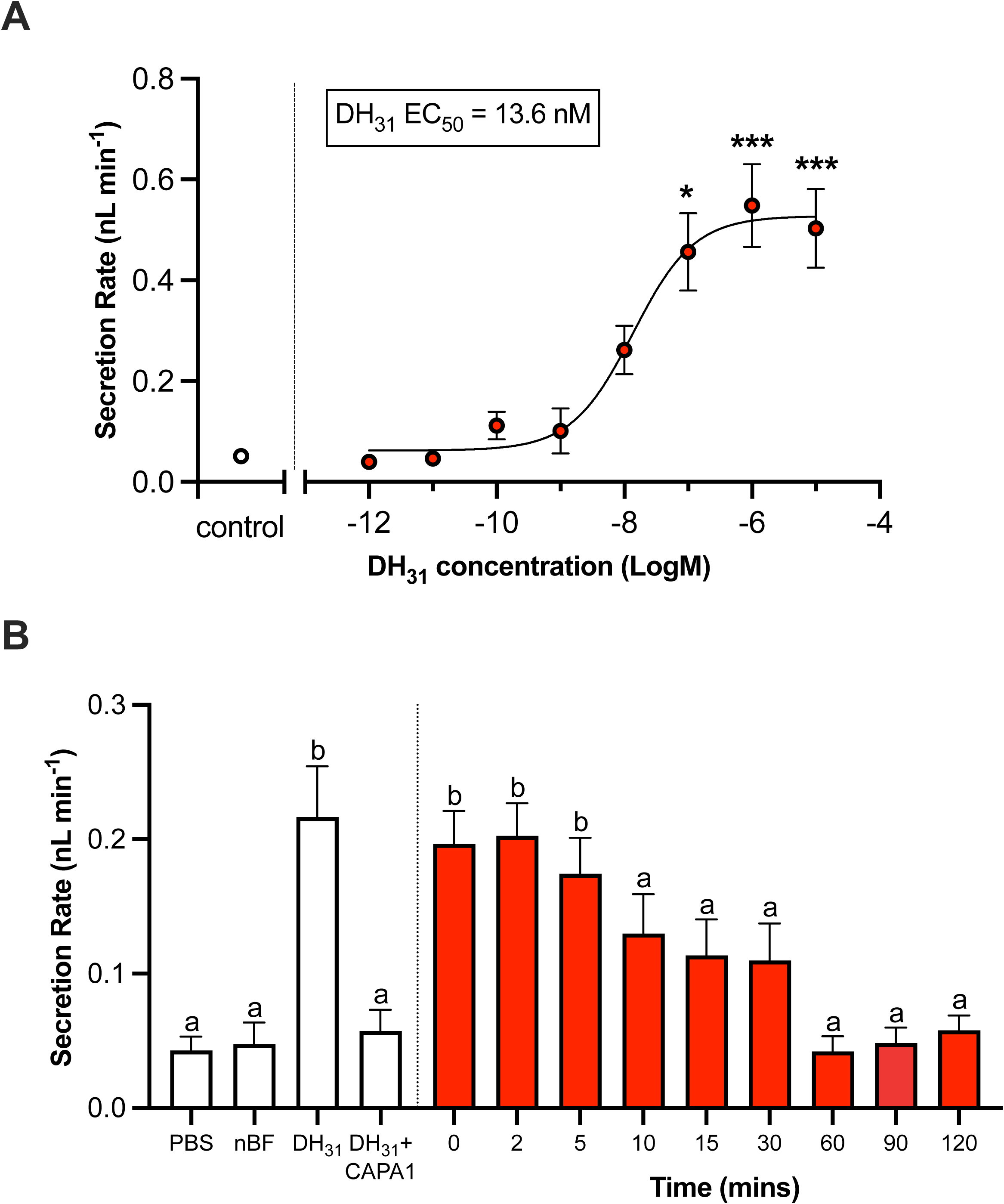
Fluid secretion by MTs from non-blood fed adult *female A. aegypti* in response to different concentrations of synthetic *Aedae*DH_31_ peptide as well as haemolymph samples from donor female mosquitoes after blood-feeding. In (A), concentrations of 10^−5^ M to 10^−12^ M *Aedae*DH_31_ were applied to isolated MTs *in vitro* for 60 min. The EC_50_ for *Aedae*DH_31_ is 13.6 nM. Significant differences between unstimulated MTs (control) and *Aedae*DH_31_-treated MTs are denoted by an asterisk as determined by one-way ANOVA with Dunnett’s multiple comparison (* denotes p < 0.01 and *** denotes p < 0.0001); data represent the mean±SEM (n=6–12). In (B), fluid secretion rates of MTs treated with diluted haemolymph samples from blood fed females post-feeding (see methods for details) for 60 min. MTs were also treated with PBS, nBF female mosquito haemolymph samples, 25 nmol l^−1^ *Aedae*DH_31_ and 25 nmol l^−1^ *Aedae*DH_31_ + 1 fmol l^−1^ *Aedae*CAPA-1 (nBF = non-blood fed). Letters above bars denote significantly different fluid secretion rates by isolated MTs treated with haemolymph samples or synthetic peptides in comparison to MTs receiving PBS alone (used as a diluent for haemolymph collected from blood-fed mosquitoes) as determined by a one-way ANOVA and Dunnett’s multiple comparison post-hoc test (p < 0.001); data represent the mean±SEM (n=9–19).

### 4.5 AedaeDH_31_ and a kinin-like peptide present synergistic activity

Considering the results above, it supports the notion that multiple diuretic hormones are released immediately post-blood feeding, playing a critical role within the first 5 minutes post-bloodmeal. To investigate potential synergistic or additive interactions between diuretic hormones, adult female MTs were stimulated with different diuretic factors alone or in combination (Fig. 5). Diuretics including DH_31_, 5HT, DH_44_, and a kinin-related peptide (CDP) significantly increase fluid secretion, while *Aedae*CAPA-1 selectively inhibits fluid secretion by DH_31_- and 5HT-stimulated MTs (Fig. 5A) as reported previously (Sajadi et al., 2018). A synergistic action (supra-additive) was observed when MTs were stimulated with both DH_31_ and CDP, since the observed secretion rate (1.15±0.23 nL min^−1^) was ∼39% higher (Fig. 5B) compared to predicted additive effects (∼0.83 nL min^−1^) calculated by summation of the mean secretion rates when DH_31_ and CDP were applied individually (Fig. 5C). The secretion rate when MTs were simultaneously treated with DH_31_ and CDP was significantly greater compared to the rate of secretion with either of these diuretics applied individually (Fig. 5C). Interestingly, while *Aedae*CAPA-1 does not affect secretion of MTs stimulated with CDP alone (Fig. 5A), the synergistic activity between CDP and DH_31_ is abolished in the presence of *Aedae*CAPA-1 (Fig. 5B) with significantly reduced secretion rates (0.34±0.18 nL min^−1^). Unlike the synergism observed by DH_31_ and CDP, no additive or synergistic activities were observed when MTs were co-stimulated with DH_31_ + 5HT or DH_44_, 5HT + DH_44_ or CDP, and DH_44_ + CDP. Similar to activity on MTs stimulated with DH_31_ + CDP, simultaneous application of *Aedae*CAPA-1 in combination with select combinations of diuretic factors, specifically DH_31_ + 5HT, DH_31_ + DH_44_ and 5HT + DH_44_, resulted in significant inhibition of fluid secretion. In contrast, *Aedae*CAPA-1 did not affect secretion rates when MTs were co-stimulated with 5HT + CDP or DH_44_ + CDP (Fig. 5B).

**Figure 5.**
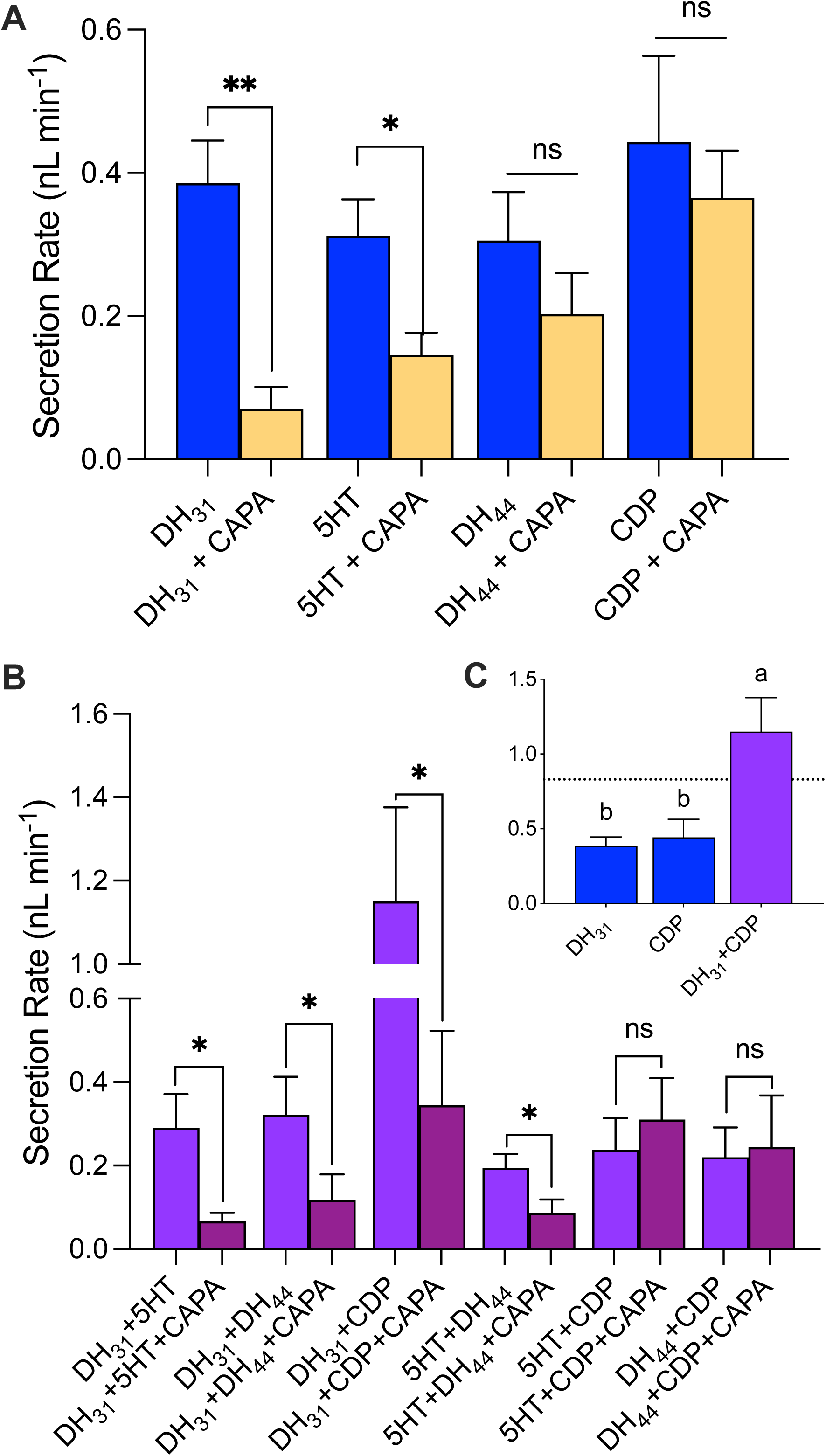
Synergistic activity between a subset of diuretic hormones and selective inhibitory activity of *Aedae*CAPA-1 on secretion rates by isolated MTs of adult female *A. aegypti.* MTs were treated with various diuretic (25 nmol l^−1^ DH_31_, 100 nmol l^−1^ 5HT, 10 nmol l^−1^ DH_44_, and/or 50 nmol l^−1^ CDP) and anti-diuretic (1 fmol l^−1^ *Aedae*CAPA-1) hormones for 60 min to test (A) individual diuretics with and without *Aedae*CAPA-1 and (B) combined actions involving two different diuretics. In (C), plot shows DH_31_ and CDP independently-induced secretion rates, along with the calculated additive effect (dashed line across y-axis at ∼0.83 nL min^−1^). In (A) and (B) main figure, significant differences between single or coupled diuretics and *Aedae*CAPA-1-treated MTs are denoted by asterisks (specifically, ** denotes p<0.001 and * denotes p<0.05) as determined by an unpaired t test (one-tailed) (mean±SEM; p<0.05, n=6–17). In (C), different letters denote significant differences in fluid secretion rates between treatments as determined by a one-way ANOVA and Tukey’s multiple comparison post-hoc test (p < 0.05); data represent the mean±SEM (n=10–17).

## 5 Discussion

In female mosquitoes, including *A. aegypti*, the evolution of blood feeding behaviour required precisely coordinating several physiological processes including digestion, excretion, and oogenesis (Klowden, 2013). While blood engorgement is necessary for egg development providing a source of nutrients, vitamins, and proteins (Beyenbach, 2003; Beyenbach and Petzel, 1987), females ingest several times their haemolymph volume in blood, which imposes an osmoregulatory challenge (Kersch and Pietrantonio, 2011). Consequently, female mosquitoes retain essential proteins from the bloodmeal in the midgut while excess water and ions are excreted from the body by the actions of the MTs and hindgut, the latter being the main reabsorptive organ (Beyenbach, 2003; O’Donnell, 2008; Williams et al., 1983). The rapid diuresis that ensues after blood gorging can result in the elimination of 40% of the ingested plasma volume within the first hour after the bloodmeal (Williams et al., 1983). Consequently, this process requires stringent control of the excretory system to ensure appropriate removal of excess ions and fluid from the body (Adams, 1999), and subsequently, modulation to this regulation to preserve nutritive elements. Multiple hormones are released into the female haemolymph to regulate diuresis, with functional genetic studies implicating roles for the CT-like (DH_31_) and kinin receptors in the post-prandial diuresis of *Aedes* mosquitoes (Kersch and Pietrantonio, 2011; Kwon et al., 2012), while recent evidence identified CAPA peptides act as critical anti-diuretic hormones (Sajadi et al., 2018; Sajadi et al., 2020; Sajadi et al., 2023). Using heterologous assays, the present study aimed to measure circulating levels of DH_31_, CAPA, and kinin peptides in the haemolymph of blood fed females to better understand the timing of hormone release yielding insight on their roles and actions post-blood feeding in *A. aegypti*.

While numerous hormones function in regulating diuresis after a bloodmeal, the mosquito natriuretic peptide, DH_31_, is of particular interest given its dual diuretic and natriuretic activity (Coast et al., 2005; Kwon et al., 2012; Sajadi et al., 2018). In mosquitoes, rapid fluid secretion during the peak phase of diuresis is driven by the action of DH_31_, acting through its receptor, DH_31_-R, increasing cAMP levels to drive secretion of Na^+^ into the tubule lumen (Coast et al., 2005; Kwon et al., 2012) facilitated by increased V-type H^+^ ATPase (proton pump) assembly and activity in principal cells (Sajadi et al., 2023). Based on current evidence in the literature, data directly confirming that the *Aedae*DH_31_-R is specifically and exclusively activated by the mosquito natriuretic (DH_31_) was lacking but was nonetheless assumed based on sequence homology when this DH_31_ receptor was molecularly identified (Kwon et al., 2012). Therefore, using a heterologous receptor assay, we determined the functional activation of *Aedae*DH_31_-R, testing various peptidergic ligands. Indeed, as expected, our findings confirm that *Aedae*DH_31_-R is specifically activated only by its natural ligand DH_31_, whereas other tested mosquito (and other insect) neuropeptides failed to activate *Aedae*DH_31_-R. Insect CT/diuretic hormone (DH) receptors belong to the secretin-receptor family of GPCRs (Hewes and Taghert, 2001) and the first insect CT/DH receptor was functionally characterized in the fruit fly, *D. melanogaster* (Johnson et al., 2005). While CT-like receptors have also been reported in other insects based on genome and protein database exploration (Caers et al., 2012), a relatively limited number have been functionally deorphanized (ie. their natural ligand identified), including in the kissing bug *R. prolixus* (Zandawala et al., 2013) and the silkworm *Bombyx mori* (Iga and Kataoka, 2015). Herein, we have functionally validated that *Aedae*DH_31_ specifically activates *Aedae*DH_31_-R and this insight was a prerequisite to reliably measure circulating levels of this hormone in the haemolymph of the female mosquito prior to and after blood feeding.

Earlier studies demonstrated that post-blood feeding in female mosquitoes, diuretic hormones rapidly induce fluid secretion within 2 minutes (Petzel et al., 1987), with secretion rates (but not the duration of diuresis) varying depending on bloodmeal size and hormone concentration (Nijhout and Carrow, 1978). Using RNA interference, it was established that *Aedae*DH_31_-R regulates fluid secretion from the MTs treated with *Aedae*DH_31_ (Kwon et al., 2012), which correlated with high excretion rates *in vivo* from female mosquitoes post-bloodmeal. Specifically, *in vitro* application of *Aedae*DH_31_ resulted in an increase in fluid secretion within 5 minutes, but this diuretic activity was blunted following knockdown of *Aedae*DH_31_-R (Kwon et al., 2012). Consistent with these previous reports, we determined that *Aedae*DH_31_ is released into the female haemolymph immediately post-blood feeding, with levels significantly elevated between 2-10 minutes (peaking at 5 minutes) while returning to levels equivalent to those in non-blood fed female mosquitoes by 15-30 minutes. In *A. aegypti* and *A. gambiae* mosquito MTs, fluid secretion increases approximately three-fold after stimulation with *Aedae*DH_31_ (Coast et al., 2005; Sajadi et al., 2018), via cAMP (Beyenbach, 2003; Coast et al., 2005), activating Na^+^ channels and the Na^+^:K^+^:2Cl^−^ co-transporter in the basolateral membrane of principal cells, subsequently activating protein kinase A (PKA), and upregulating VA activity driving cation transport processes (Dames et al., 2006; Sajadi et al., 2023; Tiburcy et al., 2013; Zimmermann et al., 2003). Notably, the EC_50_ of the endogenous natriuretic hormone (DH_31_) in *A. aegypti* to stimulate fluid secretion by adult female MTs (13.6 nM) identified in the current study aligns well with that previously reported (∼50nM) for *A. gambiae* tubules (Coast et al., 2005). *Aedae*DH_31_-R expression is found in a distal-proximal gradient, with greatest colocalization with the VA and exchangers in principal cells of the secretory distal regions of the MTs (Kwon et al., 2012; Patrick et al., 2006). The co-localization of the *Aedae*DH_31_-R, VA, and cation/proton exchangers (Kang’ethe et al., 2007; Piermarini et al., 2009; Pullikuth et al., 2006) in the secreting distal regions can explain the rapid transport of ions (mainly Na^+^), and water in response to *Aedae*DH_31_ (Kwon et al., 2012) immediately post-bloodmeal.

By heterologously expressing the *A. aegypti* kinin receptor (*Aedae*Kinin-R), we also measured the release of kinin peptides post-bloodmeal in female mosquitoes. In *A. aegypti*, three leucokinin-like peptides have been identified, *Aedes* kinins I, II, and III (Veenstra, 1994; Veenstra et al., 1997), which are released from the abdominal ganglia (Chen et al., 1994) and activate *Aedae*Kinin-R expressed in MTs and the hindgut (Kersch and Pietrantonio, 2011; Lu et al., 2011; Pietrantonio et al., 2005), to stimulate fluid secretion by the MTs and hindgut contractions (Veenstra et al., 1997) while potentially also decreasing Na^+^ reabsorption across the rectal epithelium (Lajevardi and Paluzzi, 2020), that collectively enhances fluid excretion. Previous studies have functionally characterized the *Aedae*Kinin-R, providing evidence of a single leucokinin receptor in *A. aegypti* that responds similarly (but not identically) to all three endogenous kinins (Pietrantonio et al., 2005). Through testing various peptidergic ligands, we further validated the specificity of *Aedae*Kinin-R, demonstrating that only kinin-like peptides (CDP and LK) activate this receptor with similar low nanomolar EC_50_ values as reported previously (Pietrantonio et al., 2005). The *Aedae*Kinin-R is a GPCR that activates a Cl^−^ conductance pathway via IP_3_ as a second messenger stimulating release of intracellular Ca^2+^ (Hayes et al., 1994; O’Donnell et al., 1998; Yu and Beyenbach, 2002). Unlike *Aedae*DH_31_-R localized to principal cells, the *Aedae*Kinin-R was previously immunolocalized to stellate cells of the MTs, where it functions in regulating anion permeability of the epithelium (Lu et al., 2011; Radford et al., 2002), orchestrating a switch from a moderately tight to a leaky epithelium, allowing transepithelial transport of solutes and water (Schepel et al., 2010). The current study determined that kinin-like peptides are indeed expeditiously released into the female haemolymph, peaking at 2 minutes post-bloodmeal. Relatedly, a significant decrease in excretion rate post-blood feeding was reported following *Aedae*Kinin-R knockdown (Kersch and Pietrantonio, 2011). Taken together, this indicates that *in vivo*, the *Aedae*Kinin-R is activated early during the peak phase of diuresis when kinin peptides reach their highest levels in the haemolymph and then continues in the post-peak phase. As non-selective diuretic stimulators of NaCl and KCl secretion by MTs (Beyenbach, 2003; Sajadi et al., 2018) as well as myotropic activity on the hindgut in *A. aegypti* (Veenstra et al., 1997), kinin peptides are key regulators of hydromineral balance, including roles in critical phases of the post-prandial diuresis (Kersch and Pietrantonio, 2011). Our results indicate that kinin peptides appear to be one of the first hormones released when blood-feeding ensues and are thus critical for the short-lived peak diuresis allowing the female to rid the excess salts and water derived from the bloodmeal.

Haemolymph from female *A. aegypti* had the greatest diuretic activity when samples were collected within the first 5 minutes post-bloodmeal, with lower diuretic activity observed when MTs were treated with haemolymph samples collected 10-30 minutes post-blood feeding. The decrease in diuretic activity soon after blood feeding may be due to the release of another factor into the haemolymph, initiating a feedback mechanism to inhibit cAMP signaling (Cabrero et al., 2002) leading to reduced primary urine formation by the MTs. While it had been earlier assumed that diuresis in blood feeding insects was terminated by reducing the concentration of diuretic hormones circulating in the haemolymph, (Maddrell, 1964), later studies raised the prospect that the cessation of diuresis may involve the release and actions of anti-diuretic hormones, including CAPA-related neuropeptides (Paluzzi and Orchard, 2006; Quinlan and O’Donnell, 1998; Quinlan et al., 1997). To advance our understanding of the kinetics of hormone release associated with diuresis post-blood feeding in the female *A. aegypti* mosquito, we explored whether CAPA neuropeptides are released after blood feeding in line with their established anti-diuretic activity (Sajadi et al., 2018; Sajadi et al., 2020; Sajadi et al., 2023). Indeed, CAPA peptide levels in the haemolymph are significantly increased at 30 minutes after blood feeding. Recent reports in the adult female *A. aegypti* have indicated that the two endogenous CAPA peptides bind and similarly activate their cognate receptor expressed in principal cells of the MTs (Sajadi et al., 2020), and reduce DH_31_-induced diuresis (Sajadi et al., 2018) by inhibiting VA activity and assembly in the apical membrane of principal cells (Sajadi et al., 2023). In *D. melanogaster*, CAPA peptides act as diuretic hormones on MTs isolated *in vitro* at higher doses (Davies et al., 2013; Kean et al., 2002; Terhzaz et al., 2012), but are anti-diuretic at lower concentrations (MacMillan et al., 2018; Rodan et al., 2012). Quantification of CAPA levels in the haemolymph of *D. melanogaster* indicated that resting titres are likely subnanomolar (MacMillan et al., 2018). Consistent with these earlier observations in another dipteran, the current findings revealed that circulating levels of CAPA peptides in *A. aegypti* range within picomolar concentrations. The combined action of increased CAPA levels with a reduction of DH_31_ and kinin peptides in the female haemolymph may therefore explain the decreased secretory activity observed 10 minutes after blood feeding. Thus, these dynamic hormone profiles are likely essential for functional integration whereby they coordinate the transition between the different phases of post-blood feeding diuresis described in the *A. aegypti* mosquito (Williams et al., 1983). The early peak phase is dominated by diuretic hormones including *Aedae*DH_31_ and kinin peptides, which were quantified herein and found to be significantly elevated between 2 and 10 minutes after blood-feeding. Comparatively, the post-peak phase of diuresis still involves pronounced rates of fluid secretion but is marked as a transitional stage where the excretory system switches from handling the bulk of the NaCl-rich plasma to the KCl-rich waste derived from nutritive cellular portion of the blood meal occurring during the post-peak diuresis (Williams et al., 1983).

Our results support that at least two classes of diuretic hormones, DH_31_ and kinin-related peptides, are active in the early stages of the peak phase of diuresis in *A. aegypti* since significantly elevated fluid secretion rate was evident in MTs treated with haemolymph extracts collected immediately following blood feeding. While the increase in fluid secretion mimics the immediate release of DH_31_ into the haemolymph, it further suggests the actions of additional diuretics released in concert with DH_31_. This possibility is supported by the observation that peak secretory activity was already observed in haemolymph samples collected immediately post-bloodmeal, whereas levels of DH_31_ in the haemolymph peaked at 5 minutes post-blood feeding. Earlier reports implied the potential for synergism between DH_31_ and kinin-related peptide on adult mosquito MTs, since 1µM application of each peptide individually resulted in secretion rates of 1.25-1.5 nL/min whereas in combination resulted in secretion rates ∼6 nL/min that is greater than their additive effects (Coast et al., 2005); however, this was not confirmed nor discussed in this earlier study, and the concentrations used were supraphysiological in light of our current findings. In the present study, we established that *in vitro* application of physiologically relevant concentrations of both DH_31_ and a kinin-like peptides, CDP, elicit synergistic activity on the MTs, resulting in fluid secretion rates significantly greater than their individual activities and higher than their predicted additive effects. Interestingly, addition of *Aedae*CAPA-1 together with both of these diuretic hormones abolishes the synergism observed, whereas fluid secretion rate is unaltered when *Aedae*CAPA-1 is treated with CDP-stimulated MTs alone, as reported previously (Sajadi et al., 2018). In *R. prolixus*, two diuretic hormones, 5HT and *Rhopr*DH (a CRF-related diuretic hormone), act synergistically with an observed fluid secretion rate significantly greater than their expected additive effects (Paluzzi et al., 2012b). While *Rhopr*CAPA-α2 did not influence *Rhopr*DH-stimulated secretion in isolation, addition of *Rhopr*CAPA-α2 together with both diuretic hormones (5HT and *Rhopr*DH) abolished the synergism observed (Paluzzi et al., 2012b). Comparatively, the data collected herein for *A. aegypti* indicate that DH_31_ and kinin jointly drive the rapid diuresis eliminating Na^+^-rich urine, while release of CAPA peptides into the haemolymph abolishes this action.

The female *A. aegypti* mosquito ingests a bloodmeal for the purpose of gaining proteins and other nutrients essential for egg production (Beyenbach, 2003; Beyenbach and Petzel, 1987; Christophers, 1960). However, the bloodmeal also contains large amounts of unwanted salts and water which poses a threat to the haemolymph homeostasis (Beyenbach, 2003) and reduces the maneuverability and flight speed of the mosquito (Roitberg et al., 2003). While considerable studies have examined the roles of diuretic and anti-diuretic hormones post-blood feeding, here we further functionally characterized the DH_31_-R and Kinin-R in the *A. aegypti* mosquito and provided novel evidence demonstrating DH_31_, kinin, and CAPA peptides are differentially released into the haemolymph post-bloodmeal. Based on the heterologous functional receptor assay and *in vitro* fluid secretion assays on isolated MTs, DH_31_ in the haemolymph was found in the nanomolar range and is released immediately post-bloodmeal increasing by over 100-fold to stimulate fluid secretion and reduce the Na^+^ load from the blood plasma. Accompanying DH_31_, we also provide evidence that kinin peptides are released after blood feeding, increasing over 15-fold in the haemolymph. Importantly, these two peptide hormones were found to have synergistic activity on the MTs, increasing the rate of fluid secretion beyond their individual or additive activities. Lastly, we confirmed the anti-diuretic hormone (CAPA) is released 15-30 minutes post-bloodmeal at picomolar ranges, increasing over 12-fold over basal levels, which together with decreased diuretic hormone release into the haemolymph, results in a steady decline in DH_31_-mediated natriuresis and kinin-mediated diuresis that defines the early (peak) phase of diuresis post-blood feeding, thus preventing the excessive loss of Na^+^. Taken together, these novel findings highlight the physiologically relevant titres of diuretic and anti-diuretic hormones released post-bloodmeal in the mosquito, emphasizing the fine-tuning of the excretory system by multiple hormonal factors to maintain haemolymph homeostasis.

## Author contributions

J.-P.P. and F.S. designed research; F.S., C.D., and L.S performed experiments and collected data; F.S. and J.-P.P. analyzed data and wrote the manuscript.

## Funding acknowledgement

This research was funded by the Natural Sciences and Engineering Research Council of Canada (NSERC) Discovery Grant (JPP), Ontario Ministry of Research Innovation Early Researcher Award (JPP), and NSERC CGS-D (FS) and the Carswell Scholarships in the Faculty of Science, York University (FS) along with a NSERC Undergraduate Student Research Award (CD).

